# Construction of a genetic linkage map in *Pyropia yezoensis* (Bangiales, Rhodophyta) and QTL analysis of several economic traits of blades

**DOI:** 10.1101/486175

**Authors:** Linbin Huang, Xinghong Yan

## Abstract

*Pyropia yezoensis* is an important economic seaweed, to construct a genetic linkage map and analyze the quantitative trait loci (QTLs) of blades, a doubled haploid (DH) population containing 148 DH strains established from the intraspecific hybridization between two strains with different colors was used in the present study and genotyped using 79 pairs of polymorphic sequence-related amplified polymorphism (SRAP) markers labeled with 5'-HEX and capillary electrophoresis. A chi-square test for significance of deviations from the expected ratio (1:1) on the loci which were polymorphic between parents and segregated in mapping population identified 301 loci with normal segregation (*P* ≥ 0.01) and 96 loci (24.18%) with low-level skewed segregation (0.001 ≤ *P* < 0.01). The map was constructed using JoinMap software after a total of 92 loci were assembled into three linkage groups. The map spanned 557.36 cM covering 93.71% of the estimated genome, with a mean interlocus space of 6.23 cM. Kolmogorov-Smirnov test (α=5%) of the marker positions along each LG showed a uniform distribution. After that, 10 QTLs associated with five economic traits of blades were detected, among which one QTL was for length, one for width, two for fresh weight, two for specific growth rate of length and four for specific growth rate of fresh weight. These QTLs could explain 2.29-7.87% of the trait variations, indicating that their effects were all minor. The results will serve as a framework for future marker-assisted breeding in *P. yezoensis.*

## Introduction

*Pyropia yezoensis* is a marine red alga with high nutritional values and is one of the most important maricultural crops across the world, mainly in Japan, Korea and China [1]. During the cultivation of *P. yezoensis*, hundreds of tons of nutrients (nitrogen and phosphorus) are removed by blade harvest from the eutrophic seawaters every year [2]. However, some problems such as germplasm degeneration, frequent diseases and bad harvests [3-5] have arisen under the influence of global warming [6]. Therefore, new varieties with higher yield, stronger resistant to abiotic stress and greater ecological adaptability were urgently needed for the stable development of *Pyropia* industry.

The traditional breeding methods of *P. yezoensis* are based on either observed variations by selecting blades with induced variants [3, 7, 8], or controlled crosses by selecting blades presenting recombination of desirable genes from different parents [9-11]. However, traditional breeding is usually time-consuming and less efficient [12], and has a limited ability to breed complex characters [13]. Fortunately, progress in molecular genetics has enabled plant breeders the direct selection of genotypes, thereby accelerates crop improvement [14]. Molecular marker-assisted selection (MAS) has become the main direction of plant breeding [15-17], during which construction of a genetic linkage map is one of the most important steps [18]. To date, genetic linkage maps in dozens of different species of plants and animals are successively constructed and have played important roles in various studies [19]. However, genetic linkage map construction in seaweeds was still in its infancy and was only reported in five important species, including *Laminaria japonica* [20], *L. longissima* [21], *Ectocarpus siliculosus* [22], *Porphyra haitanensis* [23] and *Undaria pinnatifida* [24]. Those maps were used for quantitative trait locus (QTL) detection of economic traits [25, 26], mapping of sex-linked loci [24, 27] and large-scale assembly of genome sequence [22]. The reasons for the relative lag of seaweed maps included the lack of polymorphic molecular markers, such as the most common used SSR markers [21, 23] and the difficulty of mapping population establishment when the parents were highly heterozygous [20]. For *P. yezoensis* in this study, the blades were monoecious and could be self-fertilized, the heterozygote could only be identified through F_1_ blades if they were mainly color-sectored [28], which depended on tissue culture techniques. There was no other mapping population of *P. yezoensis* reported except for our previous work on a doubled haploid (DH) population [29], which might be the main reason that no genetic linkage map for *P. yezoensis* was constructed.

The economically important traits of gametophytic blades of *P. yezoensis* are quantitatively inherited traits controlled by multiple genes as reported in our previous work [29]. By means of genetic mapping, quantitative traits can be decomposed into multiple QTLs, and the genetic basis of complex quantitative traits can be clarified [30]. In the present study, a genetic linkage map of *P. yezoensis* was constructed using sequence-related amplified polymorphism (SRAP) markers and high-performance capillary electrophoresis analysis based on a DH population, for further QTL detection of economic traits of gametophytic blades. The results will help application of MAS in breeding varieties for *P. yezoensis* in future.

## Materials and methods

### Plant materials

Two parental strains of *P. yezoensis* with different economic traits were used in this study. Py-HT was a red-type pigmentation mutant whose gametophytic blade had fast growth rate, thin thickness, high content of major photosynthetic pigments and high-temperature resistance. Py-LS was a wild-type strain whose gametophytic blade had slow growth rate, thick thickness, low content of major photosynthetic pigments, and poor heat-resistance [31]. In our previous work, Py-HT was used as maternal and Py-LS was used as paternal in an intraspecific cross because they were monoecious and could be self-fertilized, the heterozygote (heterozygous conchocelis) was then identified according to the methods described in Yan and Aruga (28), for the construction of a DH mapping population [29], which was then used in the present work. Briefly, only four-color sectored mosaic blades were screened from the F_1_ blades which developed from the conchospores released from the heterozygous conchocelis. Every blade was then cut into four color-sectors according to boundaries of adjacent color-sectors and every color-sector was cultured singly subsequently. A DH strain was declared when one of the carpospores was released from a self-fertilized color-sector and developed into a single conchocelis. Finally, a mapping population of 152 DH strains were established from 37 four-color sectored mosaic blades. All strains were conserved in our laboratory in the form of free-living conchocelis at 19±1 °C under a photon flux density of 10±1 μmol photons m^-2^ s^-2^ (10:14 LD) provided by cool-white, 40-W fluorescent lamps according to the method described in Kato and Aruga (32).

### DNA extraction

Genomic DNA was isolated from 30-40 mg fresh weight of free-living conchocelis of each DH and parent using a Plant Genomic DNA Kit (DP305, TIANGEN) with a modification in sample treatment. Briefly, conchocelis was sucked dry of culture solution and cut into a smooth paste in 100 μL deionized water with a single edge razor blade, instead of liquid nitrogen grounding, and then DNA was extracted according to the manufacturer’s protocol. The concentration and purity of the DNA solution were determined based on the spectrophotometric absorbance and the ratio of OD260/OD280 (Nanodrop 2000, Thermo Fisher Scientific). DNA size and integrity were assessed by 1.0% agarose gel electrophoresis. DNA with high quality was diluted to 30 ng · μL^-1^ with Tris-EDTA buffer solution and stored at -20°C for further experiments.

### Polymorphic primers screening

The sequences of 21 forward primers and 21 reverse primers (Table 1) were obtained from original papers [33-35] and designed according to the method as described in Li and Quiros (36). After random pairing, 441 primer combinations were obtained. Primers were synthesized in Sangon Biotech (Shanghai) Co., Ltd. and amplified in two parents and four DH strains to screen primer combinations with rich polymorphic loci. The PCR reaction was carried out in 15.0 μL solution containing 7.5 μL Taq PCR Master Mix (B639293, Sangon Biotech), 1.0 μL forward and 1.0 μL reverse primer (20.0 μM), 1.0 μL genomic DNA (30.0 ng · μL^-1^) and 4.5 μL deionized water. The SRAP procedure was performed as previously described in Li and Quiros (36). PCR products were separated by electrophoresis on an 8% non-denaturing polyacrylamide gel [Acryl/Bis (29:1), 1×TBE] (native-PAGE) and photographed with Gel Imaging System (Gel Doc XR+, Bio-Rad) after a rapid and economic silver staining method [37]. Briefly, gel was washed twice with deionized water for 60 s every time (the same below) and stained with 300 mL silver nitrate solution (0.1% w/v) for 15-20 min, then the gel was washed twice and developed in 300 mL sodium hydroxide solution (1.6% w/v, including 300 μL formalin) until the bands were clear with a blemish-free background, finally the gel was washed twice and photographed. Bands were detected and analyzed with Image Lab^TM^ Software (version 5.1, Bio-Rad) according to the instruction manual. Primer combinations that amplified abundant bands were screened and used to analyze the mapping population.

**Table 1.**
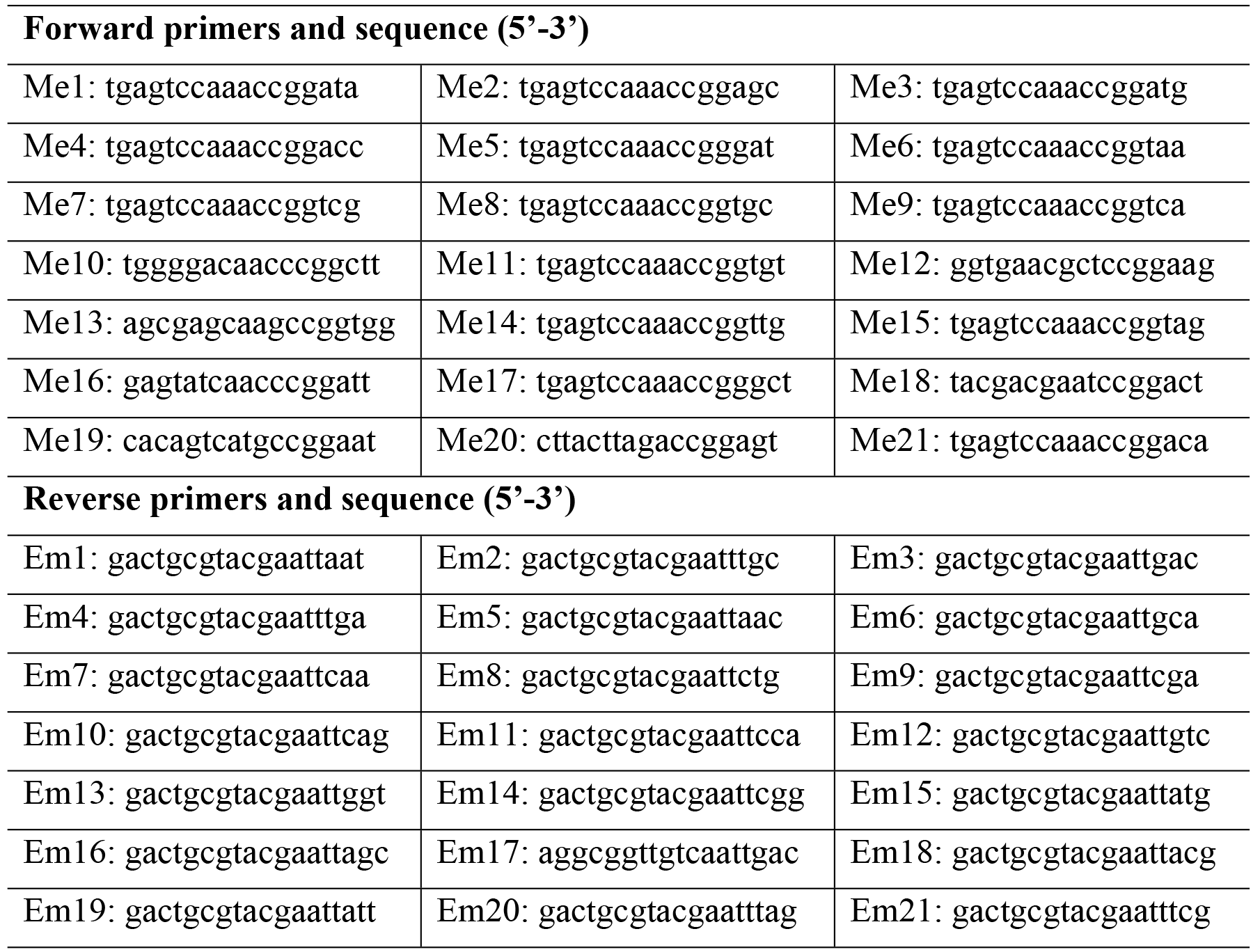
Sequence-related amplified polymorphism primers used to detect DNA polymorphisms among parents and doubled haploid population of *Pyropia yezoensis*.

### Capillary electrophoresis detection

For increased efficiency and accuracy of genotyping, the mapping population was genotyped using polymorphic primer combinations labeled with 5’-HEX (Hexachloro fluorescein phosphoramidite) under PCR conditions ibidem, and the PCR products were then sent to Sangon Biotech (Shanghai) Co., Ltd for capillary electrophoresis according to the methods as described in published studies [38, 39]. Briefly, 50 pg of amplified product was added to a mixture containing 990 μL Hi-Di formamide (Applied Biosystems) and 10 μL internal lane standard (GS1200LIZ, Applied Biosystems) after quantification, and the bands were separated using a DNA Analyzer (3730xl, Applied Biosystems) with 50-cm capillaries filled with POP-7 separation matrix (Applied Biosystems) [38]. Capillary array systems from Applied Biosystems (Foster City, CA, USA) is one of the most common for sequencing and fragment analysis [40]. Data were collected using Data Collection software (version 4.0, Applied Biosystems) and 150 FSA file were obtained for the mapping population with 148 DH and two parents after amplified by one of the primer combinations.

### Fragment analysis and genotyping

The corresponding 150 FSA files were then analyzed together using GeneMarker program (version 2.7.1, Softgenetics, LLC) under the analysis type ‘Fragment (plant)’ [40, 41]. The peak detection threshold was set at 200 RFU and the fragment size was set to be between 100-1,000 bp. GS1200LIZ size standard was used as an internal lane size standard which enabled automated data analysis, and was also essential for achieving high run to run precision in sizing DNA fragments [42]. An Excel document was exported after each analysis when report style ‘Bin Table (AFLP/MLPA)’ was selected.

### Genetic linkage map construction

The amplified results of every primer combination were analyzed and only loci which were polymorphic between parents and segregated among 148 DH population were selected for linkage analysis, using the widely used JoinMap program (version 4.0, Kyazma B.V.) [43]. Loci data were first transformed into the formats of JoinMap.

Briefly, loci identical with the maternal parent Py-HT were manually recorded as ‘a’, those identical with the paternal parent Py-LS were recorded as ‘b’, and the missing loci were recorded as ‘-’. Population type was selected as ‘DH1’. Genotype frequency of each loci was first calculated and loci with too many missing data were excluded, before chi-square (χ^2^) test was performed to determine if the genotypic frequency at each locus was deviated from the expected 1:1 segregation ratio. Normal segregation was considered as *P* values ≥ 0.01 and low-level skewed segregation was considered as 0.001 ≤ *P* < 0.01, severely skewed segregation loci whose *P* values < 0.001 were not used for linkage analysis.

The process of constructing a genetic linkage map involves a stepwise approach [44, 45]. In the present study, markers with *P* values ≥ 0.001 were first divided to different linkage groups (LGs) using command ‘Create groups using the grouping tree’, markers within each group were then ordered under the major criteria of a maximum recombination fraction of 0.4 and a minimum LOD score of 1.0 using ‘Regression Mapping’ method [46], distance between the markers were finally calculated using Haldane’s mapping function [47]. QTL IciMapping program (version 4.1, CAAS) was used to output the graphical presentation of the genetic linkage map [48].

SRAP loci were labelled according to the primer combination employed and their estimated fragment length, e.g. ‘M19E9-180.6’ designated a locus that yielded a 180.6 bp fragment with the primer combination of Me19 and Em9 (Table 1). The name of a skewed marker was suffixed by ‘D’, for example, ‘M13E2-473.6D’. Therefore, the positions of distorted markers can be easily observed from the genetic linkage map. If they were clustered in special regions on chromosomes and then these regions are termed as segregation distortion regions (SDRs) [49-51]. The presence of an SDR in the present study was declared when two or more distorted markers were clustered. The directions of distortion were determined by comparing the information of each locus with parental genotypes.

### Genome length and map coverage

The expected size of *P. yezoensis* genome (*L*) was estimated using two different methods, in which *L*_1_ is calculated as the summed length of all LGs added two times of the average marker spacing [52], and *L*_2_ is calculated as the length of each LG multiplied by the factor (m+1)/(m-1), where ‘m’ is the number of markers on each LG [53]. The estimated *L* was the average of the lengths calculated by the above two methods. Map coverage was estimated by the ratio between the cumulative map length and the expected genome size.

### Marker distribution

Marker distribution along each LG was evaluated by comparing the difference between the expected positions of the markers and the observed ones using Kolmogorov-Smirnov test (α=5%) as described in Lombard and Delourme (54). Online package KS-test was used to calculate the corresponding *D* value and *P* value of the test [55]. A random distribution of markers on each LG was indicated when *D* < *D* (Ni, 0.05) or *P* > 0.05. The values of *D* (Ni, 0.05) were determined according to Jerrold (56).

### QTL mapping

Phenotypic values of six economic traits of F_1_ gametophytic blades of the mapping population were described in our previous work [29]. These traits were blade length, width and fresh weight at the 50th day (L50, W50 and FW50), and the specific growth rate of length, width and fresh weight between 40th and 50th day (LGR, WGR and FWGR) (S1 Table). QTL was analyzed with QTL IciMapping program (version 4.1, CAAS) using the ICIM-ADD method (inclusive composite interval mapping of additive and dominant QTL) [48, 57, 58]. A stringent LOD threshold 1.5 was set to identify the putative presence of QTLs associated with economic traits of blades. A QTL was declared when the LOD value was higher than threshold of 1.5 [59]. QTLs were named and shown in italic by prepending a lower-case ‘q’ to the abbreviation of a trait name, followed by the serial number of LGs where the QTL was found and a terminal number providing a unique number to distinguish multiple QTLs of one trait on a single chromosome [60, 61], e.g. ‘*qL50-2-1*’ designated the first QTL of L50 detected on LG2.

## Results

### Genotyping

After PCR amplification, native-PAGE detection and gel image analysis, 116 SRAP primer combinations which amplified abundant polymorphic bands among the parents and four randomly selected DH strains were preliminarily screened from the 441 primer combinations. The frequencies of four forward primers (Me4, Me7, Me13 and Me19) and three reverse primers (Em6, Em8 and Em10) were found high among the 116 primer combinations. To save costs, the above seven primers were synthesized and labeled with 5’-HEX. Fluorescein-labeled primers were paired with ordinary primers before genotyping the mapping population. The amplification results of each primer combinations were analyzed in GeneMarker. An Excel document including the information of amplified bands (e.g. size and peak height) was obtained for every primer combination (Table 2). Capillary electrophoresis obtained directly the digitized information of the fragments, which was more efficient than PAGE method [62].

**Table 2.** Partial digitized information of sequence-related amplified polymorphism fragments of six samples analyzed using GeneMarker software based on the results of capillary electrophoresis.

| Sample name | Peak height (RFU) of amplified fragment with different size (bp) |  |  |  |  |  |  |  |
| --- | --- | --- | --- | --- | --- | --- | --- | --- |
|  | 106.4 | 107.4 | 139.9 | 140.8 | 175.9 | 176.9 | 188.0 | 189.0 |
| **108-HT** | - | 13500 | - | - | - | - | 3132 | 2424 |
| **108-LS** | 4199 | 4849 | - | - | 2137 | 3167 | 2365 | 2759 |
| **108-031** | 8630 | - | 1782 | 1860 | - | - | 1171 | - |
| **108-032** | - | - | - | - | - | - | - | - |
| **108-033** | - | 18946 | 38962 | 24087 | 1020 | - | - | - |
| **108-034** | - | - | 16119 | 10153 | - | - | - | - |

‘-’ denoted that no fragment whose peak height $\geq 200$ RFU was amplified.

During fragment analyzing, size calling of some samples was failed probably because of PCR or electrophoresis failure, the genotypes of these samples were then used as missing data in linkage analysis. In this study, if one primer combination missed data of more than four DH strains or of one parental strain, then the data of the primer combination would not be used for map construction. Finally, PCR data of 79 primer combinations were used for genetic mapping. As shown in Table 3, a total of 42,049 loci were amplified, of which 5,661 loci were amplified in parents with 5,172 polymorphic loci (91.36%). In addition, there were 5,059 loci amplified in both parents and the mapping population, of which 4,570 (90.33%) were polymorphic between parents and were segregated among 148 DH population.

**Table 3.**
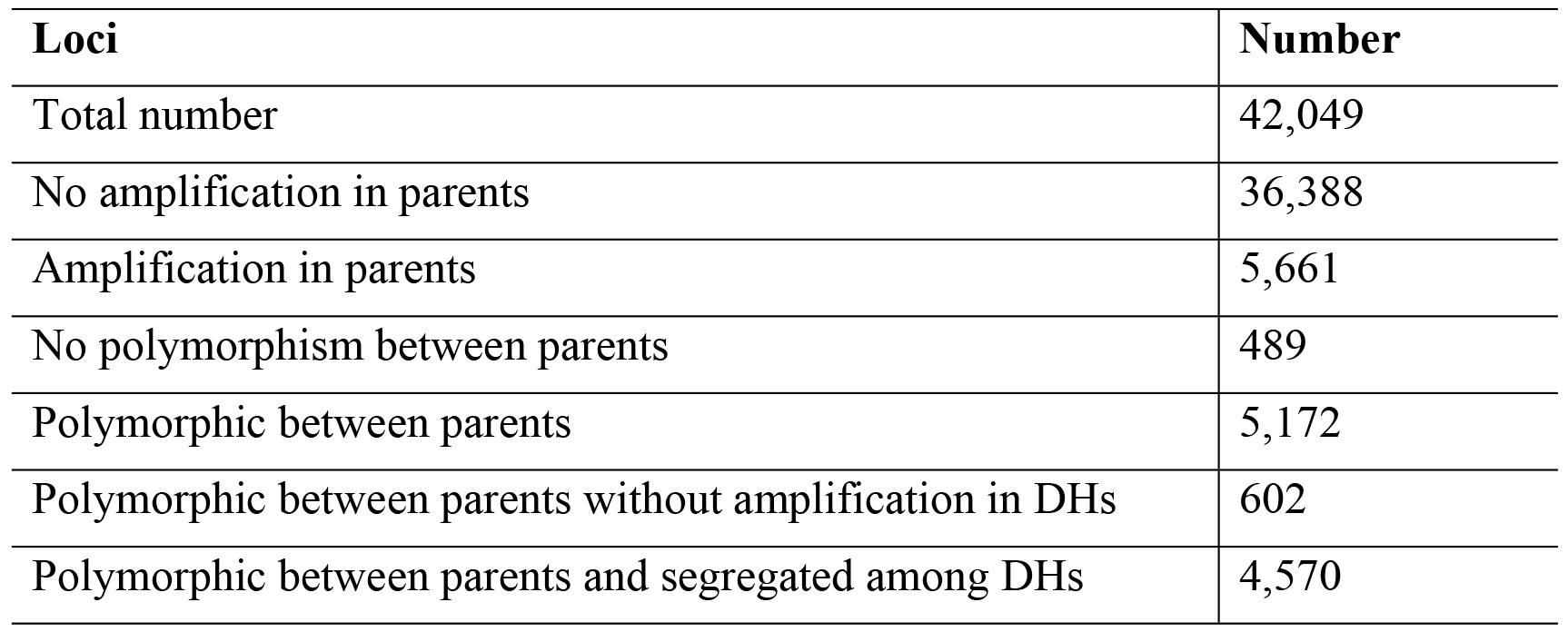
Amplified loci in parents and doubled haploid (DH) population of *Pyropia yezoensis* analyzed by fluorescent sequence-related amplified polymorphism markers.

### Map construction

A total of 4,570 SRAP loci meeting the requirement of linkage analysis were imported into JoinMap and evaluated with χ^2^ test. It was found that 301 loci were segregated with expected 1:1 ratio at *P* ≥ 0.01, and 96 loci were low-level skewed at *P* values between 0.001~0.01 (Table 4). Meanwhile, 3,775 loci with serious segregation distortion (*P* < 0.001) were discarded. Ultimately, a total of 397 loci including 227 loci specific to maternal parent Py-HT and 170 loci specific to paternal parent Py-LS were used for linkage analysis (Table 4). Approximately, five informative loci were amplified by each primer combination.

**Table 4.** Segregation of sequence-related amplified polymorphism loci in the doubled haploid population of *Pyropia yezoensis.*

| Segregation type | *P* value | $\chi^2$ value | Number of markers | Loci specific to Py-HT | Loci specific to Py-LS |
| --- | --- | --- | --- | --- | --- |
| Normal | $\geq 0.01$ | $\leq 7.81$ | 301 (75.82%) | 169 | 132 |
| Low-level distorted | 0.001~0.01 | $\leq 12.08$ | 96 (24.18%) | 58 | 38 |

At LOD 7.0, two groups with the most numbers of markers (118 and 94, respectively) were used to construct LG1 and LG2, respectively. The remaining markers were moved to a new group and those with LOD values above 4.0 were used to construct LG3. Finally, genetic linkage map of *P. yezoensis* containing three LGs was constructed (Fig 1). The map including 92 SRAP markers spanned a total distance of 557.36 cM, with a mean interlocus space of 6.23 cM between adjacent markers (Table 5).

**Fig 1.**
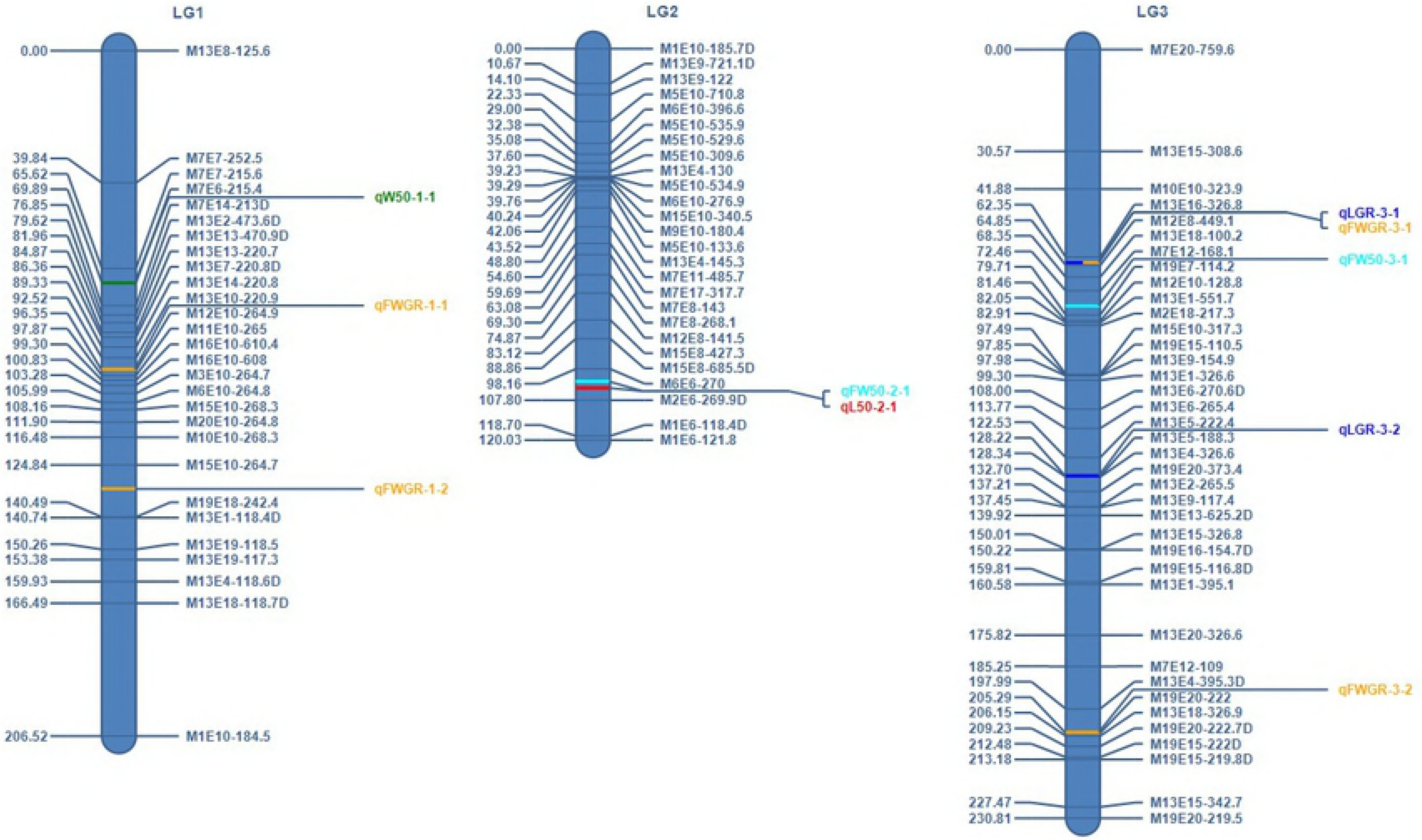
Distribution of quantitative trait loci (QTLs) controlling five economic traits of gametophytic blades on a genetic linkage map of *Pyropia yezoensis* constructed with sequence-related amplified polymorphism markers based on a doubled haploid population. The designations on the right are marker names, on the left are mapped distances in centimorgans based on Haldane’s mapping function. The colored bars denote QTLs positions and the names of QTLs are next to the long lines on the right. Loci showing low-level segregation distortion (0.001 < *P* < 0.01) are indicated with a letter ‘D’ suffix.

**Table 5.** Information on the genetic linkage map of *Pyropia yezoensis* constructed with sequence-related amplified polymorphism markers based on a doubled haploid population.

| ID | Number of markers | Length (cM) | Interlocus space (cM) |  |  | Number of gaps > 20 cM |
| --- | --- | --- | --- | --- | --- | --- |
|  |  |  | Mean | Min. | Max. |  |
| **LG1** | 28 (7) | 206.52 | 7.65 | 0.25 | 40.03 | 3 |
| **LG2** | 26 (5) | 120.03 | 4.80 | 0.06 | 10.90 | 0 |
| **LG3** | 38 (8) | 230.81 | 6.24 | 0.12 | 30.57 | 2 |

Figures in parentheses denoted the number of low-level segregation distortion markers

In addition, maximum marker spacing on the three LGs were between M13E18-118.7D and M1E10-184.5 (40.03 cM), M2E6-269.9D and M1E6-118.4D (10.90 cM), M7E20-759.6 and M13E15-308.6 (30.57 cM), respectively. The minimum spacing were between M19E18-242.4 and M13E1-118.4D (0.25 cM), M13E4-130 and M6E10-534.9 (0.06 cM), M13E5-188.3 and M13E4-326.6 (0.12 cM), respectively. Furthermore, there were five gaps large than 20.0 cM, including three gaps large than 30.0 cM and one gap larger than 40.0 cM. On a SNP-based linage map of *L. japonica*, the largest gap was 14.97 cM although the mean interlocus space was 0.36 cM [27]. Those large gaps might be due to the lack of enough polymorphisms on some chromosome regions between mapping parents [63]. Large numbers of additional genetic markers [64, 65] or mapping population from different cross combinations [66] are needed to fill in the gaps to provide a higher-resolution map.

### Genome coverage and marker distribution

Based on the total data set of the mapping population, the average estimate of expected genome length was 597.74 cM, thus the map covered 93.71% of the estimated genome of *P. yezoensis.* Furthermore, the results of Kolmogorov-Smirnov test showed that *D* values of LG1, LG2 and LG3 were 0.21, 0.19 and 0.18, respectively, all of which were smaller than the corresponding values of *D* (Ni, 0.05) (0.25, 0.26 and 0.22, respectively). The corresponding *P* values of the three LGs were 0.49, 0.67 and 0.50, respectively, all of which were larger than 0.05. These results indicated that the markers along the three LGs were in uniform distribution, which was an important feature of a high quality genetic linkage map [54].

### Segregation distortion

There were six SDRs each composed of 2-3 clustered distorted markers which had the same skew directions after being compared with the genotypes of the parents. The loci within the SDRs on LG1 and LG2 skewed toward Py-HT and Py-LS, respectively, and the loci within the first and second SDR on LG3 skewed toward Py-LS and Py-HT, respectively (Table 6). Besides, there were six isolated segregation distortion loci, of which M13E7-220.8D, M13E1-118.4D, M13E13-625.2D and M13E4-395.3D skewed toward Py-HT, and M15E8-685.5D and M13E6-270.6D skewed toward Py-LS. Genes related to segregation distortion may exist in these regions [67, 68], further study is needed to explore it.

**Table 6.** Information of segregation distortion regions (SDRs) on the genetic linkage map of *Pyropia yezoensis*.

| ID | Location | Segregation distortion marker | Skew direction |
| --- | --- | --- | --- |
| **SDR1** | LG1 | M13E13-470.9D, M7E14-213D, M13E2-473.6D | Py-HT |
| **SDR2** | LG1 | M13E18-118.7D, M13E4-118.6D | Py-HT |
| **SDR3** | LG2 | M1E10-185.7D, M13E9-721.1D | Py-LS |
| **SDR4** | LG2 | M2E6-269.9D, M1E6-118.4D | Py-LS |
| **SDR5** | LG3 | M19E15-116.8D, M19E16-154.7D | Py-LS |
| **SDR6** | LG3 | M19E15-222D, M19E20-222.7D, M19E15-219.8D | Py-HT |

### QTL mapping

In general, marker distance should be less than 10.0 cM for a map used for QTL analysis [69-71], which was 6.7 cM on the first map with a coverage of 82.8% for QTL mapping of frond length and width in *L. japonica* [26].The present map covered 93.71% of the genome and the SRAP markers along each LG were evenly distributed with an average distance of 6.23 cM between adjacent markers, which was then used for QTL detection of blade traits in *P. yezoensis.* In total, 10 QTLs associated with L50, W50, FW50, LGR and FWGR were identified (Fig 1 and Table 7). QTL associated with WGR was not detected. One QTL for L50 and one QTL for W50 were identified on LG2 and LG1, with phenotypic variance explained (PVE) of 5.72 and 7.05%, respectively. Two QTLs for FW50 were identified on LG2 and LG3, with PVE of 4.84 and 6.45%, respectively. Two QTLs for LGR were identified on LG3, with PVE of 7.87 and 3.23%, respectively. Four QTLs for FWGR were identified on LG1 and LG3 with PVE ranging from 2.29 to 4.53%.

**Table 7.** Quantitative trait loci (QTLs) of five economic traits of gametophytic blades of *Pyropia yezoensis.*

| QTL | Position<br>(cM) | Left marker | Right marker | LOD<br>values | PVE<br>(%) | Add<br>values | Interval of<br>confidence (cM) |  |
| --- | --- | --- | --- | --- | --- | --- | --- | --- |
|  |  |  |  |  |  |  | Left | Right |
| *qW50-1-1* | 70.00 | M7E6-215.4 | M7E14-213D | 1.7 | 7.05 | 0.04 | 67.50 | 74.50 |
| *qFWGR-1-1* | 96.00 | M13E10-220.9 | M12E10-264.9 | 1.8 | 2.35 | 1.16 | 92.50 | 97.50 |
| *qFWGR-1-2* | 132.00 | M15E10-264.7 | M19E18-242.4 | 1.7 | 3.20 | 1.35 | 125.50 | 140.50 |
| *qFW50-2-1* | 102.00 | M6E6-270 | M2E6-269.9D | 2.3 | 4.84 | 2.49 | 87.50 | 107.50 |
| *qL50-2-1* | 104.00 | M6E6-270 | M2E6-269.9D | 2.0 | 5.72 | 0.83 | 91.50 | 107.50 |
| *qLGR-3-1* | 64.00 | M13E16-326.8 | M12E8-449.1 | 2.1 | 7.87 | 0.90 | 63.50 | 66.50 |
| *qFWGR-3-1* | 64.00 | M13E16-326.8 | M12E8-449.1 | 1.6 | 4.53 | 1.61 | 62.50 | 66.50 |
| *qFW50-3-1* | 77.00 | M7E12-168.1 | M19E7-114.2 | 1.9 | 6.45 | -2.83 | 73.50 | 80.50 |
| *qLGR-3-2* | 128.00 | M13E5-222.4 | M13E5-188.3 | 1.6 | 3.23 | 0.59 | 122.50 | 128.50 |
| *qFWGR-3-2* | 205.00 | M13E4-395.3D | M19E20-222 | 1.7 | 2.29 | -1.17 | 201.50 | 205.50 |

PVE (%) denotes phenotypic variation explained by a QTL, Add denotes estimated additive effect of a QTL

According to published studies, PVE of major QTLs should be larger than 15% [72-74], or larger than 10% [75]. PVE of QTLs in the present study were between 2.29-7.87%, therefore they all had minor effects. Generally, only loci with major effects would undergo map-based cloning and be thoroughly studied [76]. In recent years, more attention has been given to QTLs with relatively small effects and it has shown that minor-effect QTLs also make important contributions [77]. However, they may have inconsistent additive effects across different genetic backgrounds and environments, selection of reliable candidates for further study remains a challenge [77].

LOD values of the 10 QTLs ranged from 1.6-2.3, among which three values were greater than or equal to 2.0 (Table 7). It is not clear what the threshold LOD values for QTL detection should be to declare significance [78], but LOD scores of 2.0-3.0 are commonly used to control the probability of overall false positives within 0.05 [25, 79, 80]. Higher LOD values could better control the occurrence of false QTLs and is suitable for fine mapping of major QTLs [81, 82]. However, true QTLs with minor genetic effects are not easy to detect at higher LOD values and thus LOD values may be lowered in order to detect a greater number of minor QTLs for marker-assisted breeding [80]. Therefore, these QTLs with different LOD values in the current study could be used for different researches.

Eight of the 10 QTLs had positive values of additive effect (Table 7), indicating that their favorable alleles originate from maternal parent, and the remaining two QTLs had negative Add values, indicating that the favorable alleles originate from paternal parent [83, 84]. The result was in accordance with the fact that most characters of maternal parent Py-HT were superior to paternal parent Py-LS [31]. Additive effect occurs when two or more genes source a single contribution to the final phenotype [85].

Furthermore, according to Table 7, the interval of confidence (IC) of seven QTLs ranged from 3.0-7.0 cM, IC of other three QTLs ranged from 15.0-20.0 cM. The larger the confidence interval for a QTL, the more genes may be involved and it is difficult to determine whether the QTL is composed of single gene with large effect or multiple genes with smaller effect. ICs of 15.0-20.0 cM was too large for position cloning [86] and should be narrowed down for the precise estimate of QTL position [87, 88]. The Complex Trait Consortium considered that the interval should less than 1.0-5.0 cM for fine mapping [89], therefore, four QTLs (*qFWGR-1-1, qLGR-3-1, qFWGR-3-1* and *qFWGR-3-2*) in the present study met the requirement and could be used for further study.

Two clusters of QTLs were detected on two of the three LGs (Fig 1). The first cluster contained two QTLs (*qL50-2-1* and *qFW50-2-1*) which were 2.00 cM apart and located on LG2 within the 98.16-107.80 cM region. The distance was 3.80 cM between *qL50-2-1* and the nearest marker M2E6-269.9D and was 3.84 cM between *qFW50-2-1* and its nearest marker M6E6-270. IC of *qFW50-2-1* covered that of *qL50-2-1.* The second cluster contained two QTLs (*qLGR-3-1* and *qFWGR-3-1*) located on LG3 within the 62.35-64.85 cM region at the same position. The distance between these two QTLs and the nearest marker M12E8-449.1 was 0.85 cM. The IC of *qFWGR-3-1* covered that of *qLGR-3-1.* Mohan, Nair (90) considered that a marker should co-segregate or be closely linked with the desired trait, and 1.0 cM or less is probably enough for MAS. Therefore, six QTLs in the present study which were less than 1.0 cM distant from the nearest markers could be used for further study.

## Discussion

Historically, color mutants are used as genetic markers for crossbreeding and genetic study of *P. yezoensis* [9, 28]. The distances between the centromeres and the loci of four color mutants have been determined and assigned to three different LGs [91], which could be considered as a traditional genetic linkage map, however, the information was far from further utilization because of too little markers. In the present study, for the first time, we constructed a genetic linkage map of *P. yezoensis* containing 92 polymorphic SRAP markers and 10 QTLs associated with economic characters of blades based on a DH population. The map will provide a foundation for molecular breeding in *P. yezoensis.*

Mapping population is the basic of linkage analysis and is usually obtained from controlled cross, in which the crossing parents should have sufficient variation for traits of interest at both DNA and phenotypic level [92]. Theoretically, the higher the variation, the easier to obtain abundant recombination. However, parents should not be so diverse that they are unable to cross [93]. Our previous study revealed that the crossing parents (Py-HT and Py-LS) had significant differences in blade traits and several recombinant strains had been screened [31]. Besides, we found that the genetic similarity index between the parents was 0.4962, which suggested a high genetic diversity between them [94]. In the present study, 91.36% polymorphic loci were found between the parents after SRAP analyzing. Therefore, DH population was constructed based on the cross of Py-HT × Py-LS [29] and then used for the genetic mapping. A DH is a genotype formed when a haploid cell undergo induced or spontaneous chromosome doubling [95]. DH can be exploited to produce completely homozygous lines, to construct genetic linkage maps, to locate genes of economic importance and to increase breeding efficiency [96]. DH is especially powerful for analyzing quantitative traits because replicated traits can be analyzed re-using identical genetic material [97].

In the case of *P. yezoensis*, the blades were monoecious and could be self-fertilized, so the procedure of DH development could be accomplished by self-fertilization, this is because spermatia (male gamete) and carpogonium (female gamete) always occur diffusely on a single sector [91, 98, 99], which was developed from one of the tetrad cells after mitosis [91]. Therefore, the gametes formed on a single color-sector were genetically identical. So, we considered that self-fertilization of a color-sector was equal to the procedure of chromosome doubling of gamete. DH population of 148 strains was constructed using 37 four-color sectored F_1_ gametophytic blades [29]. Every strain of the DH population was obtained from one single color-sector and was developed from one carpospore after self-fertilization of the sector. Therefore, a DH of *P. yezoensis* was the same with that of higher plant, which were all homozygous diploid.

Based on simulation studies, the type and size of experimental population can exert an influence on the accuracy of a genetic linkage map [100]. The higher the number of individuals, the more precise is the map, but at the same time larger population means excessive work and costs associated with phenotyping and genotyping [101, 102]. It is important to select an appropriate size of population. Most experiments have used a total of 100 to 200 individuals or progenies [25, 54, 103]. Therefore, 148 DH strains was used in the present study, which is of similar size (157 strains) to *P. haitanensis* in a similar study [23].

Molecular markers are important tools for creating a genetic linkage map and have provided a significant increase in genetic knowledge in many cultivated plant species [14]. SRAP markers used in the present study is a PCR marker system which combines simplicity, reliability and a moderate throughput ratio [36]. It has been extensively used in genetic diversity analysis [104, 105] and genetic mapping in economic plants [14, 36, 106, 107], including one of the most important seaweeds in China, *P. haitanensis*, when there are not enough SSR markers [23]. SRAP can also be useful for QTL mapping because of their ability to target gene-rich regions of the genome [14]. The quantitative trait data could then be used to determine if any SRAP markers are closely associated with those traits [108]. Once the markers are identified, breeders can select desirable QTLs without interference from environmental effects [109].

During the construction of a molecular genetic linkage map, the most difficult and complicated steps are the separation of PCR products and detection of polymorphic bands. The traditional method used to separate PCR products and detect polymorphic bands is PAGE [36]. However, PAGE cannot give the accurate size of DNA fragments and it has a low detection efficiency. Besides, the method has some degree of error when the results were manually recorded. In addition, our previous study found that 11 SRAP primer combinations amplified 95.42% polymorphic bands in six strains of *P. yezoensis*, with an average of 11.4 loci per primer combination [94]. The abundant polymorphism met one of the requirements of a molecular marker for genetic mapping, but in the meantime added difficulties to the artificial recording of bands. To solve this problem, capillary electrophoresis with fluorescence detection was applied in the present study, which has advantages of high separation efficiency, short analysis time and high-throughput [110].

A phenomenon called segregation distortion which means that many markers deviate from the expected Mendelian fraction is often encountered during genetic mapping [19, 111]. Segregation distortion has been found in many studies in plants and is considered one of the main forces in biological evolutionary [112-114]. Various factors have been suggested to cause segregation distortion [115], however, the underlying mechanism is still debated and obscure [116-118]. For DH population, high percentage of segregation distortion may be caused by strong zygotic selection, that is the gametophytic competition during zygote formation [117, 119]. The percentage of skewed SRAP markers (24.18%) of the DH population in the present report was less than that (30.10%) reported previously in *P. haitanensis* [23]. Several studies show that segregation distortion affects the estimation of genetic distance and the order of markers on the same linkage group [120, 121]. Skewed markers may have some genetic information, but their accuracies are unknown, thus some researchers think they should be ignored for the accuracy of genetic linkage maps [122], as in the study of P. *haitanensis* [23]. However, if distorted markers are ignored, map coverage may decrease and some important information in the real data analysis of QTL mapping may lose [123, 124]. Several genetic linkage maps using second-generation markers contain some skewed markers [118, 125, 126]. In the present study, the genetic linkage map contained 20 skewed markers, of which 14 markers formed six SDRs, and the coverage was 93.71%, which was higher than that (88.1%) of *P. haitanensis* without skewed markers [23].

Two clusters composed of QTLs for different traits were found on LG2 and LG3 in the present study. The phenomenon of QTL cluster exists widely in crops [116, 127] and has also been found in *P. haitanensis* [25].Traits clustered within the same region are significantly correlated with each other [29, 128]. This cluster phenomenon could be considered as multifactorial linkages followed by natural selection favoring co-adapted traits and is partly due to pleiotropy of some unknown key factor(s) controlling various traits through diverse metabolic pathways [129].

## Conclusions

The SRAP genetic linkage map constructed in the present study provided a framework for linkage analysis and QTL detection in *P. yezoensis*. By saturating the map and validating these QTLs, functional markers could be identified or converted for marker-assisted breeding in future.

## Acknowledgements

This work was supported in part by the National Natural Science Foundation of China (Grant No. 31302185), the National High Technology Research & Development Program of China (‘863’ Program) (Grant No. 2012AA10A411). The funders had no role in study design, data collection and analysis, decision to publish, or preparation of the manuscript.

## Supporting information

**S1 Table. Information on the genetic linkage map and economic traits data for QTL mapping.** Sheet 1 is the marker information on the genetic linkage map, Sheet 2 is the loci information used to construction the genetic linkage map, Sheet 3 is the six economic traits data of the DH mapping population.

